# Investigating the impact of motion in the scanner on brain age predictions

**DOI:** 10.1101/2023.08.08.552504

**Authors:** Roqaie Moqadam, Mahsa Dadar, Yashar Zeighami

## Abstract

**Introduction:** Brain Age Gap (BAG) is defined as the difference between the brain’s predicted age and the chronological age of an individual. Magnetic resonance imaging (MRI)-based BAG can quantify acceleration of brain aging, and is used to infer brain health as aging and disease interact. Motion in the scanner is a common occurrence that can affect the acquired MRI data and act as a major confound in the derived models. As such, age-related changes in head motion may impact the observed age-related differences. However, the relationship between head motion and BAG as estimated by structural MRI has not been systematically examined. The aim of this study is to assess the impact of motion on voxel-based morphometry (VBM) based BAG.

**Methods:** Data were obtained from two sources: i) T1-weighted (T1w) MRIs from the Cambridge Centre for Ageing and Neuroscience (CamCAN) were used to train the brain age prediction model, and ii) T1w MRIs from the Movement-related artifacts (MR-ART) dataset were used to assess the impact of motion on BAG. MR-ART includes one motion-free and two motion-affected (one low and one high) 3D T1w MRIs. We also visually rated the motion levels of the MR-ART MRIs from 0 to 5, with 0 meaning no motion and 5 high motion levels. All images were pre-processed through a standard VBM pipeline. GM density across cortical and subcortical regions were then used to train the brain age prediction model and assess the relationship between BAG and MRI motion. Principal component analysis was used to perform dimension reduction and extract the VBM-based features. BAG was estimated by regressing out the portion of delta age explained by chronological age. Linear mixed effects models were used to investigate the relationship between BAG and motion session as well as motion severity, including participant IDs as random effects. We repeated the same analysis using cortical thickness based on FreeSurfer 7.4.1 and to compare the results for volumetric versus surface-based measures of brain morphometry.

**Results:** In contrast with the session with no induced motion, predicted delta age was significantly higher for high motion sessions 2.35 years (t = 5.17, p < 0.0001), with marginal effect for low motion sessions 0.95 years (t = 2.11, p=0.035) for VBM analysis as well as 3.46 years (t = 11.45, p < 0.0001) for high motion and 2.28 years (t = 7.54, p<0.0001) for low motion based on cortical thickness. In addition, delta age was significantly associated with motion severity as evaluated by visual rating 0.45 years per rating level (t = 4.59, p < 0.0001) for VBM analysis and 0.83 years per motion level (t = 12.89, p<0.0001) for cortical thickness analysis.

**Conclusion:** Motion in the scanner can significantly impact brain age estimates, and needs to be accounted for as a confound, particularly when studying populations that are known to have higher levels of motion in the scanner. These results have significant implications for brain age studies in aging and neurodegeneration. Based on these findings, we recommend assessment and inclusion of visual motion ratings in such studies. In cases that the visual rating proves prohibitive, we recommend the inclusion of normalized Euler number from FreeSurfer as defined in the manuscript as a covariate in the models.

## Introduction

Brain aging is a heterogeneous process characterized by both morphological and functional brain changes. These changes, which occur throughout one’s lifespan, are closely intertwined with shifts in behavior, both contributing to and being affected by it (Yankner et al., 2008). These changes are commonly observed in the aging population and are distinct from their counterparts in neurodegenerative disorders. However, aging is also the primary risk factor for most neurodegenerative disorders, including Parkinson’s disease (PD) and Alzheimer’s disease (AD) (Collier et al., 2011; Hou et al., 2019). As such, the study of aging-related brain changes has been of interest, to develop a baseline characteristic against which abnormal brain alterations can be identified (Bethlehem et al., 2022; Kaufmann et al., 2019). Similarly, at the individual level, brain aging profile has been proposed as a biomarker for brain health (Franke and Gaser, 2019).

Neuroimaging techniques have been used to study the brain structural changes throughout lifespan. In particular, T1-weighted (T1w) Magnetic resonance imaging (MRI) is widely used in research and clinical settings to study different aspects of brain morphology and its longitudinal changes in a non-invasive manner. Using MRI to investigate brain structure, function, and its abnormalities at high spatial resolution has become a prominent part of the clinical diagnostic routine (Lerch et al., 2017; Rocca et al., 2017; Rüber et al., 2018; Scheltens et al., 2021). Neuroimaging studies of normal brain aging have generated an extensive body of literature regarding the trajectories of the aging brain (Bethlehem et al., 2022; Ferreira and Busatto, 2013; Fjell et al., 2014; Lockhart and DeCarli, 2014). In addition, the need for a valid biomarker of brain aging over the lifespan has been emphasized due to the increasingly recognized importance of interventions to prevent aging related disorders such as AD (Belsky et al., 2015; Elliott et al., 2021).

Due to the individual differences in aging processes (i.e. trajectory and rates across the population as well as throughout the lifespan), biological age, that is, estimated age of a sample, and its offset in comparison with chronological age have been widely used as measures reflecting an individual’s healthy or unhealthy aging (Belsky et al., 2015; Elliott et al., 2021; Ferrucci and Kuchel, 2021; Rivero-Segura et al., 2020; Wu et al., 2021). Concerning the brain, this offset, or “Brain Age Gap” (BAG) is defined as the difference between the brain’s predicted age and the chronological age of an individual. BAG can be measured using neuroimaging data alongside machine learning models to predict the age of the participants in a healthy aging population (training data). The model is then applied to the population of interest (test data) to calculate BAG and perform group comparisons or assess relationships with phenotypes of interest. BAG can quantify acceleration of brain aging, and is used to infer brain health as aging and disease interact (Cole and Franke, 2017; Franke and Gaser, 2019). BAG has proven sensitive to various neurological and neuropsychiatric conditions, not only from the spectrum of dementia in late-life, but also in much younger patients with multiple sclerosis (Høgestøl et al., 2019), schizophrenia (Nenadić et al., 2017), obesity (Zeighami et al., 2022), first episode psychosis (Kolenic et al., 2018), diabetes (Franke et al., 2013), gestational Age (Luders et al., 2018), MCI (Franke and Gaser, 2012), traumatic brain injury (Cole et al., 2015), HIV (Cole et al., 2017c), epilepsy (Pardoe et al., 2017), and Down syndrome (Cole et al., 2017a). BAG has also been used to track brain health improvement in several intervention studies including the effect of bariatric surgery and exercise on brain health (Zeighami et al., 2022).

While in pediatric populations, age is negatively associated with presence of motion in the scanner (Baum et al., 2018; Pardoe et al., 2016), motion levels are significantly higher in the older populations (Madan, 2018; Pardoe et al., 2016; Pollak et al., 2023) as well as in individuals with obesity (Beyer et al., 2020; Zeighami et al., 2021), PD, and other disorders (Makowski et al., 2019; Torres and Denisova, 2016). Previous studies have indirectly shown that head motion is associated with alterations in T1w structural brain measure estimations. As such, head motion may also impact the observed age-related differences. Motion in the scanner is a common occurrence that can affect the acquired MRI data and act as a major confound in MRI-derived models. Madan et al. (2018) investigated the impact of head motion as measured by fMRI-estimated head motion as well as ‘average edge strength’ from T1w MRIs on various cortical characteristics and found a subtle yet significant impact of motion on cortical thickness estimates (Madan, 2018). Similarly, Savalia et al. examined the impact of head motion during fMRI scans on T1w estimated brain measures, where they found visual quality ratings (QC ratings) and FD each significantly affect cortical thickness (Savalia et al., 2017).

With regards to brain age prediction, using fMRI estimated motion, Liem et al. conducted motion regression to address excessive head motion in functional scans. While they concluded that there was minimal effect of motion, they also found that regressing out motion, as measured by framewise displacement (FD) in fMRI, significantly increased their prediction error. When they repeated their analyses on matching subjects with mean FD between 0.19 and 0.28 mm, they found no effect of motion on the performance (with an exclusion criterion of mean FD > 0.6). However, excluding cases with FD values that are greater than 0.6 might not be feasible in the older population as well as those with neurological diseases. For example, participants with Alzheimer’s dementia presented with motion levels up to 3.0 mm and above (Dai et al., 2015; Pini et al., 2020), participants with Parkinson’s disease exhibited motion in the range of 1-1.2 mm (Guo et al., 2022), and stroke patients had motion in the range of 1-2 mm which can exacerbate the effect of motion (Seto et al., 2001).

While the effect of motion has been thoroughly studied in functional MRI analysis, including the effect of motion in brain age prediction (e.g., as discussed in (Liem et al., 2017)) the relationship is not as clear when studying structural MRI. This is partially due to lack of a well-defined and agreed upon measure of motion in the T1w MRI scans. Madan et al. has used “average edge strength” from morphological analysis as a measure of motion related to cortical thickness, however, their suggestion of using fMRI estimated motion as a confound might not be practical (Madan, 2018). There are several studies which utilize Euler number as measured using FreeSurfer as exclusion criteria (De Lange et al., 2022) or include it as a covariate (Kaufmann et al., 2019) in the BAG studies. However, Euler number has been proposed as a general measure of quality and its relationship with motion has not been previously established. As such, the relationship between head motion and BAG has yet to be systematically examined (Kaufmann et al., 2019; Madan, 2018; Savalia et al., 2017; Wang et al., 2019).

The aim of this study is to address this gap and assess the impact of motion on BAG estimation based on voxel-based morphometry (VBM) as well as cortical thickness, and to evaluate whether including motion (estimated through visual assessment or an automated proxy) as a confounding factor (i.e. as estimated motion ratings) in brain age estimation models would sufficiently account for this confound and improve the accuracy and validity of the findings. Our findings can contribute to the understanding of the relationship between motion and brain age estimations and provide insights into the potential implications of motion in brain age studies, particularly in the usage of BAG in study of aging and neurodegenerative disorders.

## Methods

### 2.1. Data

Data were obtained from two different sources to train the brain age prediction model and evaluate the influence of motion on brain age predictions. These distinct sources served the following purposes: 1) training the brain age prediction model, 2) investigating the potential impact of motion on the BAG.

#### 2.1.1. Training Data

The brain age prediction model was trained using data from participants in the second stage of the Cambridge Centre for Ageing and Neuroscience (CamCAN, https://www.cam-can.org/index.php?content=dataset) dataset (Taylor et al., 2017). Participants with neurological and psychiatric conditions were excluded from the study. The training data was matched with the testing dataset and only included CamCAN participants who passed QC with ages below 50 years. A total of 281 (148 female) were included in the analysis, with age range between 18 and 50 (and mean +/- std of 36.14+/- 8.6 years). T1-weighted MRIs were acquired on a 3T Siemens TIM Trio, with a 32-channel head-coil using a 3D magnetization-prepared rapid gradient echo (MPRAGE) sequence (TR = 2250 ms, TE = 2.99 ms, TI = 900 ms, FA = 9 deg, field of view (FOV) = 256 × 240 × 192 mm, 1 mm3 isotropic, GRAPPA = 2, TA = 4 min 32 s). For detailed acquisition parameters see: https://camcan-archive.mrc-cbu.cam.ac.uk/dataaccess/pdfs/CAMCAN700_MR_params.pdf.

#### 2.1.2. Movement-related artifacts (MR-ART) dataset

The independent out-of-sample dataset used to assess the impact of motion on BAG comes from a longitudinal investigation aimed at enhancing the understanding of motion artifacts and providing a valuable resource for evaluating and improving existing motion correction techniques (Nárai et al., 2022). The dataset comprises brain scans from 148 healthy participants, the pipeline failed for one subject, we used 434 sessions from the 147 remaining subjects (7 sessions missing). Furthermore, 27 sessions failed the QC for brain mask, linear, or nonlinear registration. The remaining data comprised 127 subjects with all three sessions included (subjects with QC fails for any of the sessions were completely removed). Due to the low number of participants with ages over 50 and a gap in the age distribution in the remaining sample we also removed 6 subjects (all females) with ages over 50. The final sample includes 121 subjects, 77 females and 44 males, aged 26.17 ± 6.32 years. None of the participants had a reported history of neurological or psychiatric diseases. Brain scans were performed using a Siemens Magnetom Prisma 3T MRI scanner with a 20-channel head-neck receiver coil at the Brain Imaging Centre. T1-weighted 3D magnetization-prepared rapid gradient echo (MPRAGE) anatomical images were acquired with 2-fold in-plane GRAPPA acceleration and a 1 mm³ spatial resolution. The scans involved three conditions for each participant: a standard scan (STAND) without motion, a low head motion scan (HM1), and a high head motion scan (HM2). Participants focused on a fixation point during the scans and nodded their heads in response to the word “MOVE” on the screen for HM1 and HM2 scans.

**Table 1.** Dataset description for the training and motion related test datasets.

| Dataset | N | Age range | Sex |
| --- | --- | --- | --- |
| MR-ART | 127 (x3 = 381) | 18-67 (27.8 +/- 9.9) | Female 83 / Male 44 |
| CamCAN | 640 | 18-88 (54.2 +/- 18.61) | Female 324 / Male 316 |
| MR-ART-matched | 121 (x3 = 363) | 18-45 (26.2 +/- 6.3) | Female 77 / Male 44 |
| CamCAN-matched | 281 | 18-50 (36.2 +/- 8.64) | Female 148 / Male 133 |

### 2.2. Quality Control (QC) and Motion Assessment

#### 2.2.1. QC Protocol

For T1w data, the QC protocol involved the following main steps: assessment of presence of artifacts other than motion (e.g. missing slices, incomplete field of view), as well as linear and nonlinear alignment of the images with the outline of the MNI-ICBM152-2009c average template (Manera et al., 2020). Registration QC was conducted following the protocols outlined in Dadar et al. 2018, 2022 (Dadar et al., 2022, 2018). Each image in the two datasets underwent these steps, and quality ratings were recorded.

#### 2.2.2. Motion Rating: Visual Inspection and Rating Scale

Every image within the CAMCAN, MR-ART datasets that passed the QC steps outlined in section 2.2.1 underwent a thorough visual inspection process to assess presence and severity of motion (RM). The QC images included sagittal, coronal, and axial slices covering the entire brain, allowing for reliable visualization of motion across the entire scan (see Figure 1 for an example).

**Figure 1.**
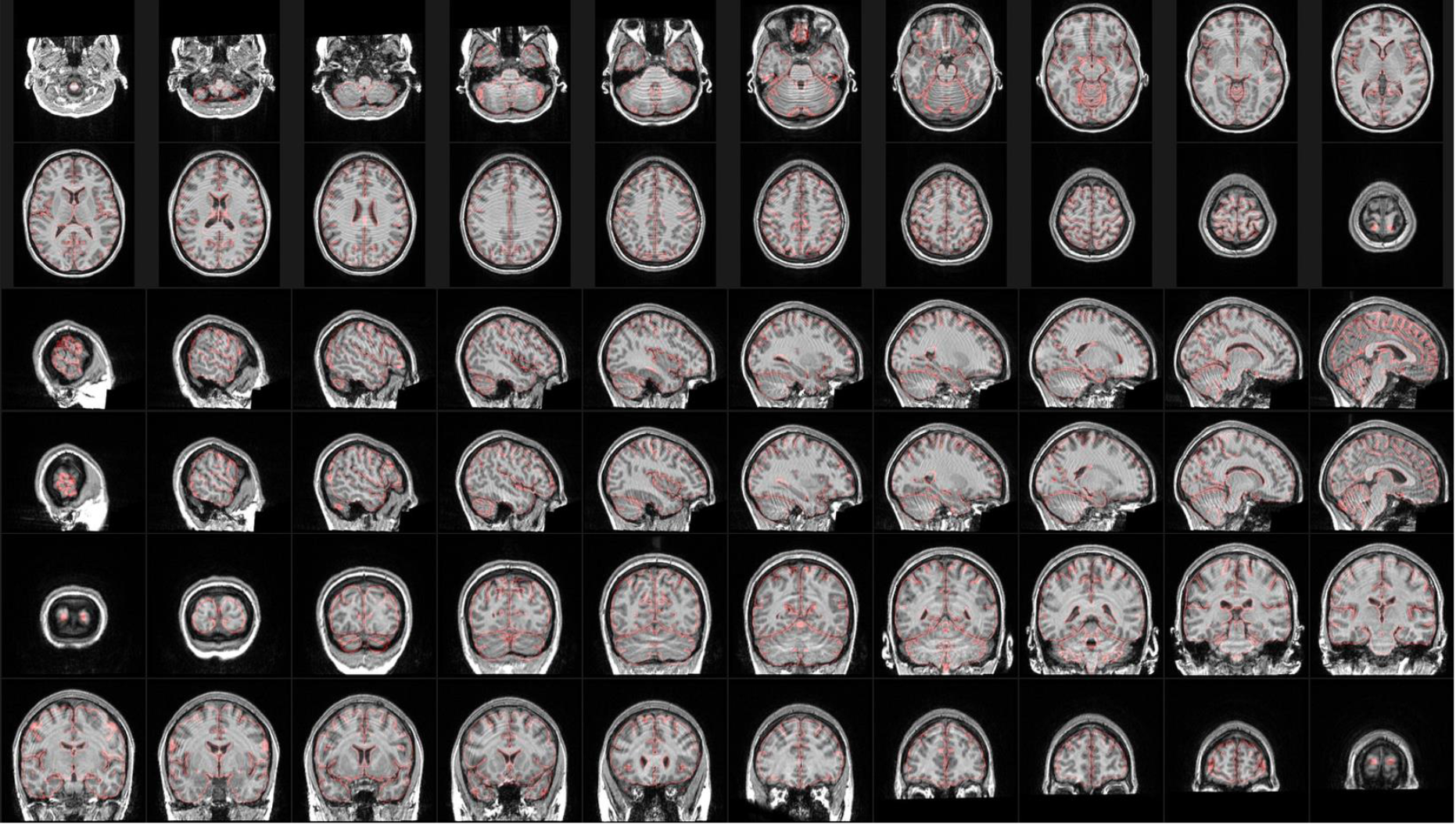
Example QC images showing sagittal, coronal, and axial slices covering the entire brain, used for visual assessment of motion severity (visual motion severity rating = 5, session HM2).

The expert rater evaluated each image, assigning a motion score based on a 6-point scale. The scale ranged from 0, indicating no motion, to 5, indicating high levels of motion. The motion level ratings were assigned in increments of 1 to provide a more detailed assessment of motion artifacts present in the images. This process ensured the accuracy and reliability of the datasets for further analysis and research purposes. Figure 2.A shows representative examples of cases with motion ratings ranging from 0-5. RM repeated the rating process a second time, 6 months after performing the initial ratings and blind to the initial scores, to allow for assessment of intra-rater reliability in motion ratings (Figure 2.B).

**Figure 2.**
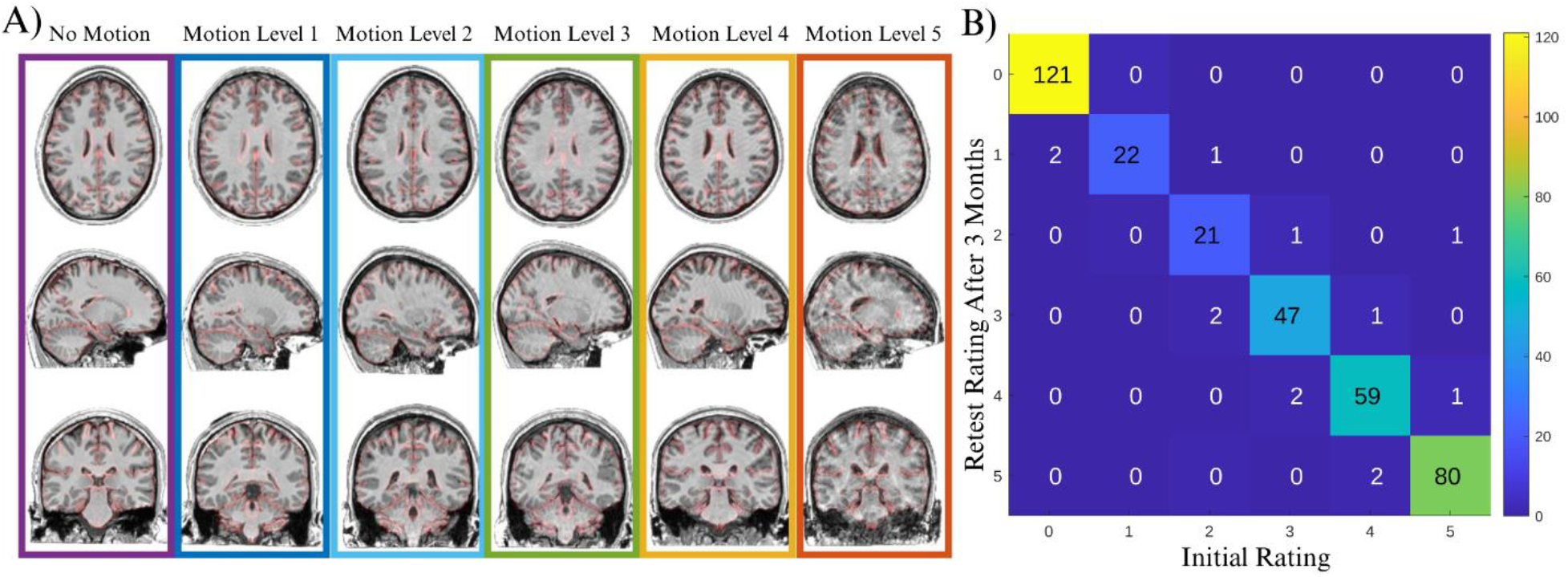
A) Representative examples of cases with no motion (rating: 0) to high motion (rating: 5) based on visual assessment. B) intra-rater agreement for visual rating of motion in MR-ART dataset.

### 2.3. Voxel-based morphometry

To train the brain age prediction model and investigate the association between predicted brain age and motion, VBM was employed to estimate gray matter density in cortical and subcortical regions. The T1w structural scans underwent a standard VBM pipeline using the following steps: 1) image denoising (Coupé et al., 2008); 2) intensity non-uniformity correction (Sled et al., 1998); and 3) image intensity normalization into range (0–100) using histogram matching. Linear registration to an average brain template (MNI ICBM152–2009c) was performed using a nine-parameter registration (Dadar et al., 2018; Manera et al., 2020), followed by a nonlinear registration using the Advanced Normalization Tools (ANTS) software (Avants et al., 2009). Segmentation of the T1w images into gray matter, white matter, and cerebrospinal fluid was performed using the ANIMAL software (Collins and Evans, 1997). Finally, VBM gray matter density maps per voxel were generated using MNI MINC tools by VBM analysis (“BIC-MNI Software repository,” n.d., p.).

### 2.4. FreeSurfer

To assess the impact of neuroimaging pipeline and features used on the results of the brain age prediction models, the analyses were repeated using cortical thickness and volumetric measurements from FreeSurfer (Fischl, 2012). Both the CAMCAN and MR-ART datasets underwent processing using FreeSurfer version 7.4.1. Raw T1 scans were reanalyzed, including subcortical segmentation and surface-based morphometry, and vertex-wise maps of cortical thickness were constructed. Cortical structure alignment between individual surfaces was achieved through spherical registration, and the resulting maps were then subjected to smoothing with a full-width at half-maximum (FWHM) of 5 mm. Furthermore, Euler numbers were extracted and employed as a quality control measure (Rosen et al., 2018), as previously described by Kaufmann et al. (2019). The Euler number serves to assess the topological complexity of the cortical surface in MRI scans, with more negative values indicating lower scan quality. The formula used to derive the Euler number is 2-2n, where n represents the number of surface holes. This computation is performed separately for the left and right hemispheres, with subsequent averaging to obtain one Euler number per participant, following the work by Rosen et al. (2018) (Rosen et al., 2018). Brain age prediction was performed using FreeSurfer-based cortical thickness.

Due to skewed distribution of the Euler number we used the following formula to transform the average Euler number for the two hemispheres to a more normalized distribution:

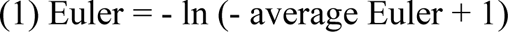

From here on we use normalized Euler numbers throughout the manuscript. The results were then compared across the three motion sessions of MR-ART data as well as 6 visual motion rating using a mixed effects model:

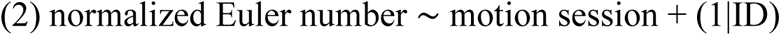

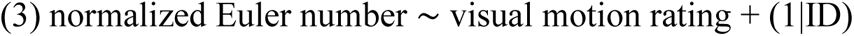

Brain age prediction was performed using FreeSurfer-based cortical thickness and subcortical volumes.

### 1.5. Brain Age Prediction Model

To predict brain age, similar to our previous works (Zeighami et al., 2022; Zeighami and Evans, 2021) and similar works in the literature (Franke and Gaser, 2019; Lewis et al., 2018; Peng et al., 2021; Smith et al., 2020, 2019), a linear regression model was employed using VBM measures as features, following principal component analysis (PCA) for dimensionality reduction. PCA is a data factorization method based on singular value decomposition that has frequently been employed in studies focused on brain age prediction. To ensure the model’s generalizability and prevent overfitting, we employed 10-fold cross-validation. The prediction accuracy obtained from the 10-fold cross-validation was utilized to evaluate the model’s performance on the training sample (i.e. CamCAN dataset). It also assisted in determining the optimal number of features to include in the final model. For the linear regression model, the root-mean-squared error (RMSE) was employed as the natural cost function. Finally, two other brain age prediction models were also trained based on the STAND (without motion) and HM2 (high motion) MR-ART sessions, to investigate i) within dataset impact of motion on brain age predictions as well as ii) the effect of using motion-impacted data for training brain age predictions models.

To further validate our brain age prediction model, we also used support vector machine regression (SVMR; with linear, gaussian, and polynomial kernels), ensemble models including Bootstrap aggregation (BAGging), as well as least absolute shrinkage and selection operator (Lasso or L1) regularization and Ridge (L2) regularization for the linear regression method to perform brain age prediction. SVMR with linear kernel and GPR showed similar performances to the linear model for the VBM analysis and only Linear Regression with Lasso regularization outperformed the linear model (Table 2). Given the better performance and interpretability of the linear regression model with regularization, we used this model throughout the analyses. For the brain age modeling based on the cortical thickness (as measured by FreeSurfer), linear regression with L2 regularization showed the next performance and therefore was used for the main analysis (Table 2). All the analyses were performed and validated with both MATLAB version 2022b and scikit-learn (Kramer, 2016) from Anaconda 2023.03. In order to make the results comparable to other works in the literature the results in Table 2 are provided based on the full CamCAN dataset. We obtained very similar performances to those reported with other methods in the literature (e.g. XG Boost by (De Lange et al., 2022), R= 0.889 and RMSE = 8.427 years in the CamCAN dataset). Similar to the work by (Han et al., 2022) comparing the performance of 27 different machine learning models (linear and nonlinear), we found the regularized linear regression algorithms achieved similar or better performance to nonlinear and ensemble algorithms.

**Table 2.**
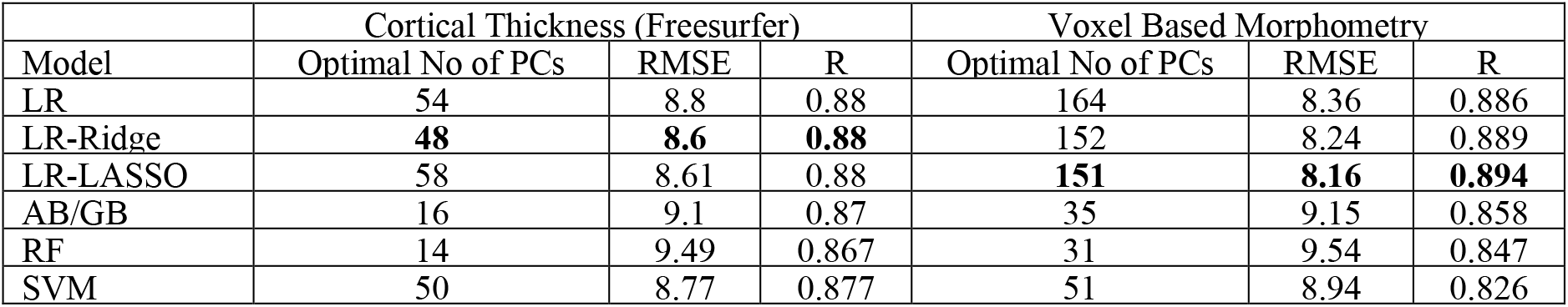
performance for FreeSurfer and VBM age prediction results (based on Scikit Learn)

|  | Cortical Thickness (Freesurfer) |  |  | Voxel Based Morphometry |  |  |
| --- | --- | --- | --- | --- | --- | --- |
| Model | Optimal No of PCs | RMSE | R | Optimal No of PCs | RMSE | R |
| LR | 54 | 8.8 | 0.88 | 164 | 8.36 | 0.886 |
| LR-Ridge | **48** | **8.6** | **0.88** | 152 | 8.24 | 0.889 |
| LR-LASSO | 58 | 8.61 | 0.88 | **151** | **8.16** | **0.894** |
| AB/GB | 16 | 9.1 | 0.87 | 35 | 9.15 | 0.858 |
| RF | 14 | 9.49 | 0.867 | 31 | 9.54 | 0.847 |
| SVM | 50 | 8.77 | 0.877 | 51 | 8.94 | 0.826 |

### 1.6. Assessing the Relationship between BAG and Motion Severity

Once the brain age was predicted as described earlier, we computed the BAG, which represents the difference between the predicted brain age and the chronological age. BAG serves as an indicator of brain-related health, as it quantifies the gap between the brain age estimated from gray matter density and the expected brain age based on chronological age.

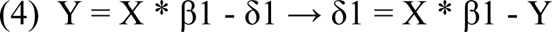

In formula (4), δ1, X, β1, and Y indicate BAG, VBM-based principal components, the weights estimated by the model, and chronological age, respectively. BAG is inherently correlated with the participants’ chronological age, which can act as a confounding factor in the analysis and complicate the differentiation between the effect of chronological age and the additional biological BAG. To address this issue, various adjustments have been proposed in previous studies (Beheshti et al., 2019; Le et al., 2018; Liang et al., 2019; Smith et al., 2019). In this study, we adopt the definition proposed by Smith et al. in 2019, where we regress out the portion of BAG that can be explained by chronological age (as shown in formula (5)) (Smith et al., 2019). The residual after accounting for the influence of chronological age is referred to as the adjusted BAG.

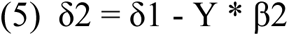

δ2 and β2 in formula (5), represent the adjusted BAG and the estimated weights by the model, respectively. The adjusted BAG serves as the primary measure of interest in this work.

### 1.7. Statistical analyses

For the MR-ART dataset, the trained models were utilized to predict brain age for the participants, not accounting for the presence of motion artifacts in this dataset. The model used for prediction was trained on the CamCAN dataset. This allowed us to assess the effect of motion artifacts on the BAG within this dataset. Following the predictions, we calculated the adjusted BAG for the test sample. Two sets of linear mixed effects models were used to assess whether presence of motion impacted the adjusted BAG estimates, one with session (contrasting the sessions with motion against the session without motion) as a categorical variable of interest contrasting the two sessions with motion (HM1 and HM2) against the session without motion (STAND), and another with the motion scores as a continuous variable of interest. All models included participant ID as a categorical random effect.

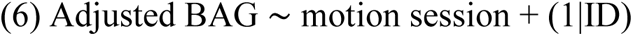

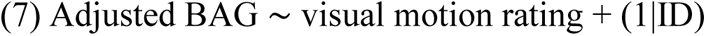

All these analyses were repeated using the normalized Euler number as follows:

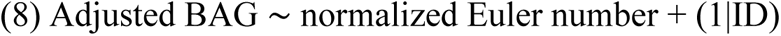

We also evaluated the effect of motion across brain age prediction features. To do so, we used the first 45 principal component (PC) maps derived from the training dataset and included in the prediction model. In order to identify the optimal number of PCs, we used 10-fold cross validation with a variable number of PCs from 1 to 200 and chose the best performing model which was then applied to the MR-ART data for the main analysis. These PCs were projected to the MR-ART data normalized based on the CamCAN dataset (as was done for brain age prediction analysis). These projections were then used in linear mixed-effect models to evaluate whether there is any main effect of motion session (both low and high session) as well as the visual motion rating on the MR-ART projection of PCs. The results were then corrected for multiple comparisons using false discovery rate (FDR) correction based on Benjamini-Hochberg.

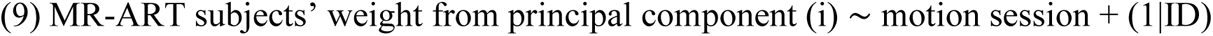

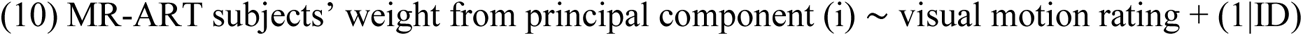

All the prediction and statistical analyses described were performed using MATLAB 2022b.

## 3. Results

### 3.1. Motion session, motion rating, and normalized Euler number

Figure 2.b summarizes the intra-rater agreement results for the visual motion ratings in the remaining MR-ART sample, showing excellent intra-rater agreement across the two rating sessions. Figure 3 shows the correspondence between MR-ART motion sessions and visual motion ratings (Figure 3.B), as well as examples of scans with different visual motion scores within the same MR-ART motion session (Figure 3.A). We further utilized Euler number as a commonly used measure reflecting the quality of the surface-based analysis to evaluate the differences across session and visual rating categories. However, given the high skewness of the Euler number (Kurtosis = 17.7), we have used the normalized Euler number as calculated by formula (1), which results in a significant reduction in kurtosis (kurtosis= 3.3). Figures 3.C and 3.D show the average normalized Euler number per subject across the three sessions as well as the 6 motion rating values.

**Figure 3.**
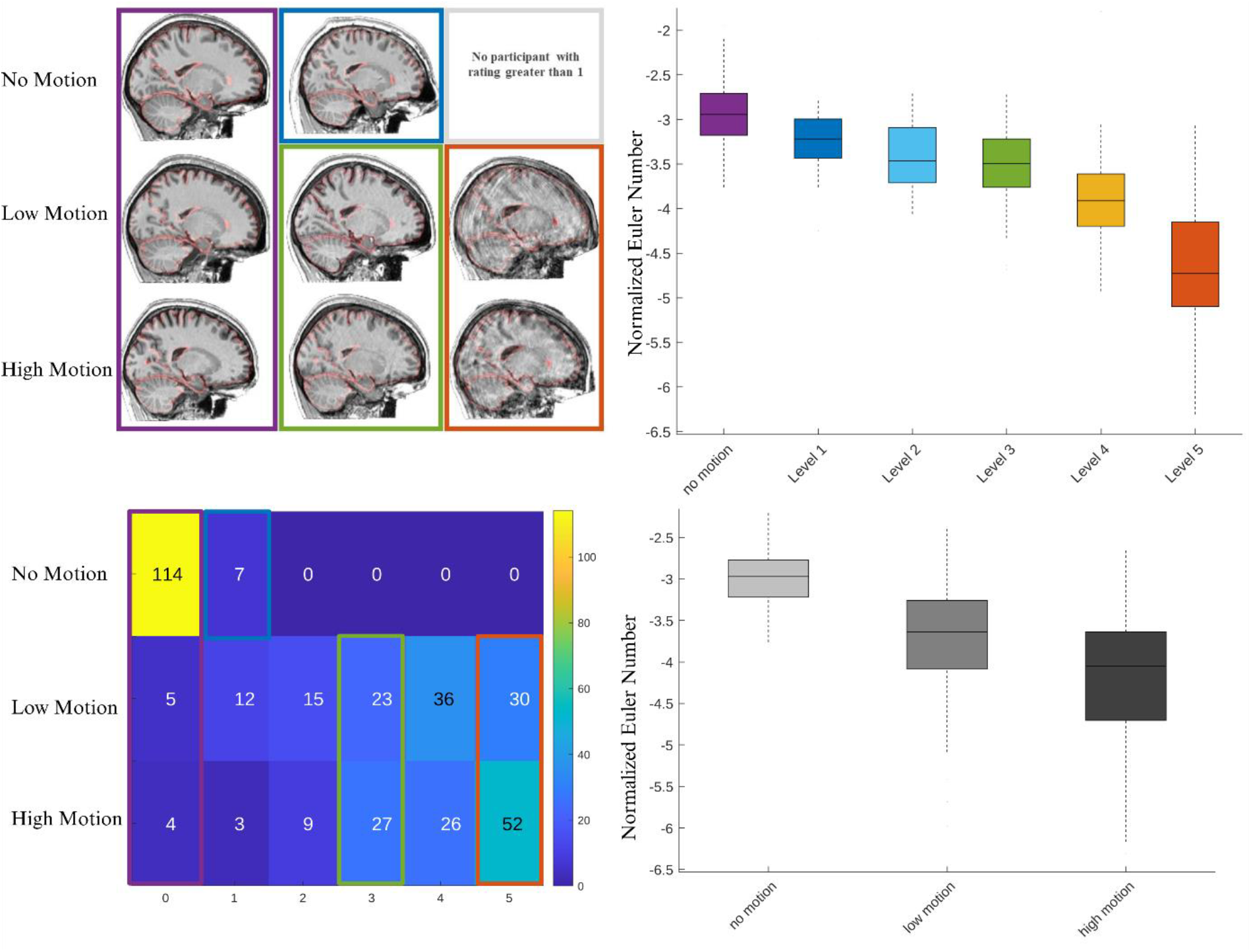
A) Correspondence between session and visual rating. Purple box shows example cases with motion rating values of 0 across the three motion sessions. Blue shows an example of a case with level 1 motion in the no-motion session (there were no higher motion ratings in the no-motion session). Green box shows examples of cases with motion level 3 in low and high motion sessions. Orange box shows examples of cases with motion level 5 in low and high motion sessions. B) Numerical values for number of overlapping cases across session and visual ratings. We have also visualized C) normalized Euler numbers as a function of visual motion rating as well as D) motion session.

Furthermore using mixed effect model, we found a significant decrease in the quality of T1w MRIS as reflected by more negative normalized Euler numbers in low motion (β = -0.77, t= -11.2, p<0.0001 for normalized Euler number; β = -36.1, t= -5.0, p<0.0001 for non-normalized Euler number) as well as high motion session (β = -1.22, t= -17.7, p<0.0001 for normalized Euler number; β = -72.5, t= -10.0, p<0.0001 for non-normalized Euler number). The model showed an adjusted R-squared of 0.59 for normalized Euler number compared to adjusted R-squared of 0.33 for non-normalized Euler number. Similarly, there was a significant effect of the overall visual motion ratings (β = -0.3, t= -24.3, p<0.0001 for normalized Euler number; β = -18.9, t= -13.2, p<0.0001 for non-normalized Euler number) with an adjusted R-squared of 0.70 for normalized Euler number compared to adjusted R-squared of 0.41 for non-normalized Euler number. These results suggest that the visual motion rating is highly related to image quality as captured by the normalized Euler number.

### 3.2. Brain age prediction model

Using the linear regression model with regularization trained based on the principal components of the gray matter VBM, it was possible to predict the chronological age of the CamCAN dataset with a correlation of r = 0.89 (p<0.0001), RMSE of 8.16 for BAG for the complete sample. Similarly for cortical thickness analysis based on FreeSurfer results, we found a correlation of r = 0.88 (p<0.0001), RMSE of 8.6 for BAG for the complete sample. As expected, the performance drops in a more limited sample with r = 0.8 (p<0.0001) based on the VBM analysis and r = 0.76 (p<0.0001) based on the cortical thickness analysis for the matched sample (The RMSE for the models go down but it is due to a more limited age range).

The age prediction model was evaluated based on the number of the principal components included in the training dataset, which is the result of 10-fold cross validation and using RMSE as the cost function for age prediction. We determined the optimal number of principal components by selecting the value that yielded the lowest RMSE. The results were N = 151 principal components for the full sample and N=45 for the matched sample for the VBM analysis and N = 48 principal components for the full sample and N=40 for the matched sample for the cortical thickness results. Consequently, our predictive models rely on 45 VBM-based principal components as the key features for prediction. In order to calculate the features for the analysis, the principal components were projected to the out-of-sample MR-ART dataset.

Boxplots of adjusted BAGs for each motion session in the MR-ART dataset are illustrated in Figure 4.A. In sessions where motion was induced, we found that the predicted delta age was significantly higher compared to sessions without induced motion. This was observed in both the low-motion and high-motion sessions, as demonstrated by the respective beta values of 0.95 years (t = 2.11, p=0.035) and 2.35 years (t = 5.17, p < 0.0001) with an adjusted R-squared of 0.49 (around 50% variance explained). These findings highlight the adverse impact of induced motion on the estimation of delta age, suggesting that motion can considerably influence the assessment of brain health. The results based on the cortical thickness analysis were similar but more pronounced as demonstrated by the beta values of 2.28 years (t = 7.54, p<0.0001) for low motion and beta of 3.46 years (t = 11.45, p < 0.0001) for high motion with an adjusted R-squared of 0.74 (around 75% variance explained).

**Figure 4.**
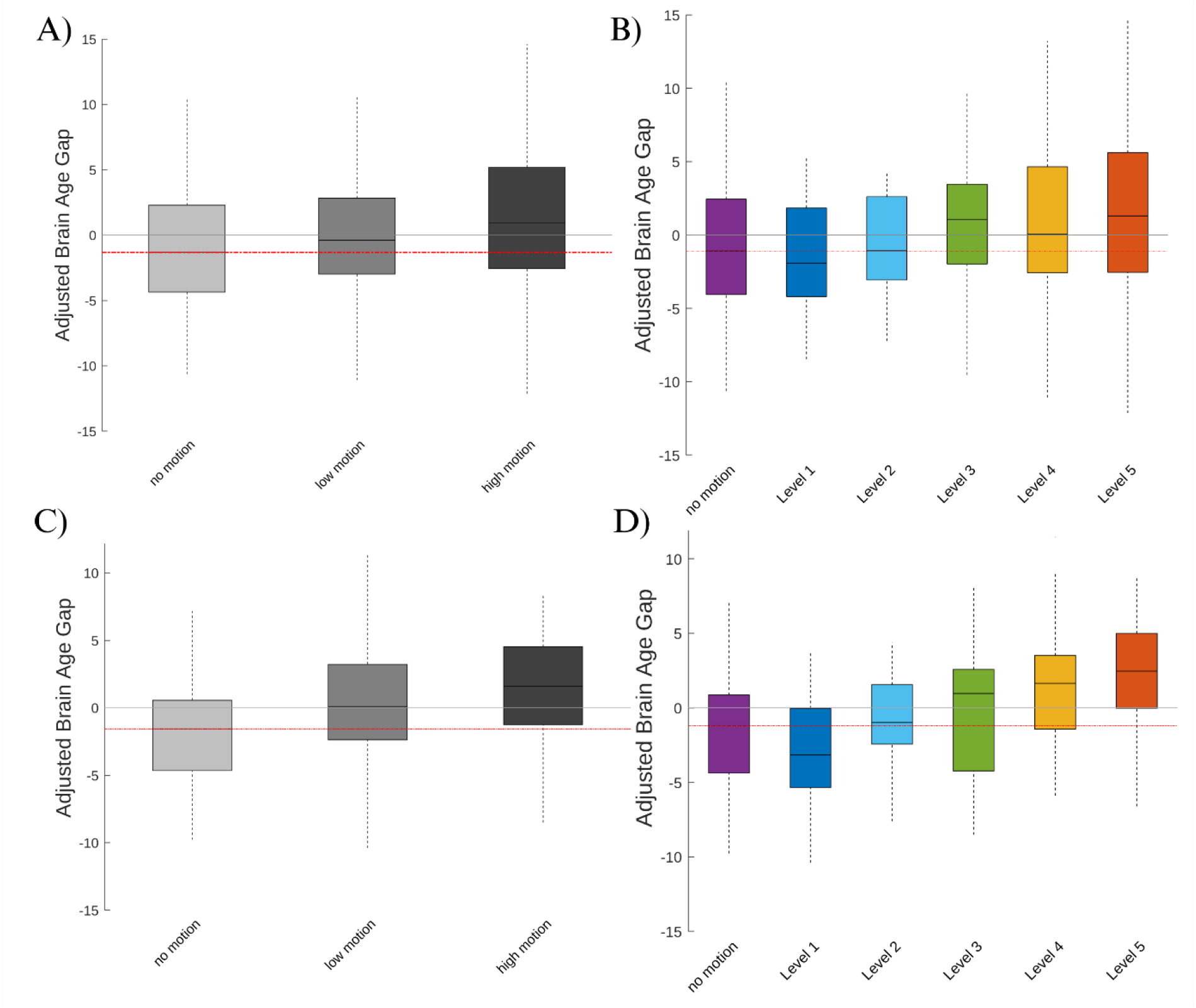
A) Adjusted brain age gap across sessions based on VBM. B) Adjusted brain age gap across visual motion rating levels based on VBM. C) Adjusted brain age gap across sessions based on cortical thickness. D) Adjusted brain age gap across visual motion rating levels based on cortical thickness.

Furthermore, we explored the relationship between motion severity (as assessed through visual ratings) and BAG. The results showed a significant association between motion severity and BAG, further highlighting the relationship between motion artifacts and the estimated age-related brain changes. Specifically, we observed that for every unit increase in motion rate as assessed by the visual rating, the estimated delta age increased by 0.45 years per rating level (t = 4.59, p < 0.0001) with an adjusted R-squared of 0.48 (around 50% variance explained) similar to the value found based on session. These findings highlight the critical importance of considering and accounting for motion artifacts when interpreting delta age estimates. Figure 4.B visually represents these findings. Similar to the analyses based on session, here again the results based on the cortical thickness analysis were similar but more pronounced as demonstrated by the beta values of 0.83 years per motion level (t = 12.89, p<0.0001) with an adjusted R-squared of 0.76 (around 75% variance explained).

Finally, we repeated the same analysis using the normalized Euler number as a proxy for motion. The results showed a significant association between normalized Euler number and BAG. Specifically, we observed that for every unit decrease in image quality based on normalized Euler number, the estimated adjusted delta age increased by 1.12 years (t = -4.2, p < 0.0001) with an adjusted R-squared of 0.47 based on the results from VBM analysis and the estimated adjusted delta age increased by 2.52 years (t = -14.7, p < 0.0001) with an adjusted R-squared of 0.80 based on the results from cortical thickness analysis. These findings highlight the possibility of using normalized Euler number as proxy of motion when studying and interpreting delta age estimates.

### 3.3. Brain age prediction model validation across sessions

Finally, as another validation step, we trained the brain age prediction model with cross-validation on the no motion session, and applied it to data from low and high motion sessions, yielding similar results. In comparison with the STAND session with no motion, there was a significant increase in the adjusted BAG in both low (t = 2.24, p = 0.025) and high (β = 0.99 years, t = 6.93, p < 0.0001) motion sessions with an adjusted R-squared of 0.9 (90% variance explained). Similarly, we observed a significant increase in the adjusted BAG for higher motion severity as assessed by the visual ratings (β = 0.19 year/rating level, t = 5.94, p < 0.0001) with an adjusted R-squared of 0.89 (approximately 90% variance explained). We also repeated this analysis a second time, training on high motion data and testing on the entire dataset. The results were very similar for adjusted BAG between no motion and high motion sessions (β = 0.79 years, t = 5.49, p < 0.0001) but no significant differences were found between no motion and low motion sessions (β = 0.15 years, t = 1.14, p > 0.05). We also found a significant relationship between adjusted BAG and the visual motion ratings (β = 0.12 year/rating level, t = 4.33, p < 0.0001). In both cases the models had an adjusted R-squared of 0.9. We found very similar results using the cortical thickness measures based on FreeSurfer analysis. Using STAND session with no motion as training data we found significant differences in both high motion (β = 1.49 years, t = 7.92, p < 0.0001) and low motion (β = 1.32 years, t = 7.02, p < 0.0001) compared to no motion session with an adjusted R-squared of 0.76. When using high motion data from FreeSurfer for training the results are very similar with significant differences in both high motion (β = 0.89 years, t = 4.9, p < 0.0001) and low motion (β = 1.06 years, t = 5.84, p < 0.0001) compared to no motion session with an adjusted R-squared of 0.76.

**Figure 5.**
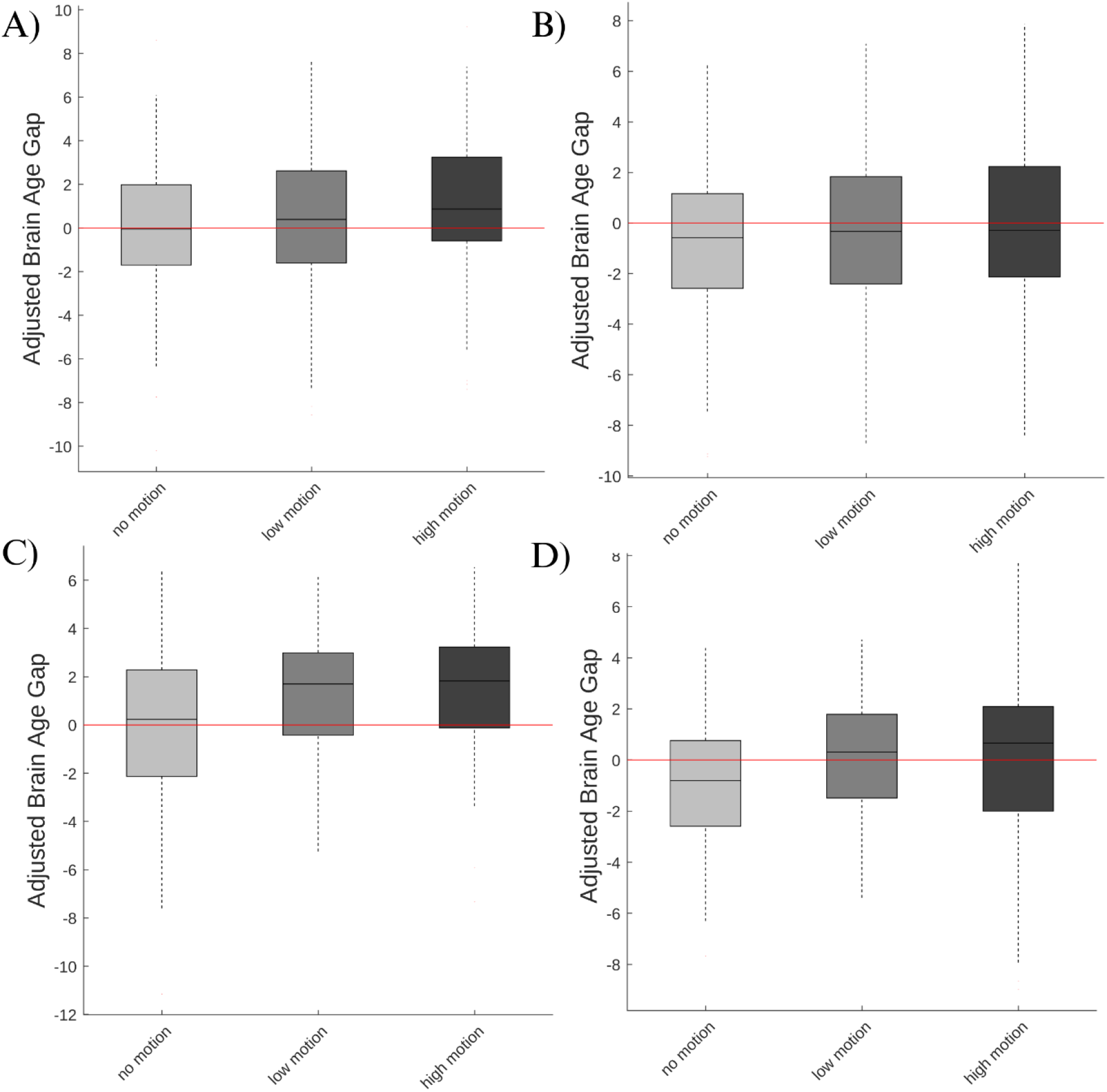
A) Adjusted brain age gap trained on the no-motion (STAND) session with cross validation, applied to all sessions based on VBM data. B) Adjusted brain age gap trained on the high-motion (HM2) session with cross validation and applied to all sessions based on VBM data. C) Adjusted brain age gap trained on the no-motion (STAND) session with cross validation, applied to all sessions based on cortical thickness data. D) Adjusted brain age gap trained on the high-motion (HM2) session with cross validation and applied to all sessions based on cortical thickness data.

### 3.4. Effect of motion on features

In order to evaluate the effect of motion across brain age prediction features, we ran similar mixed-effect models to evaluate whether there is any main effect of motion session (both low and high session) as well as the visual motion rating on the predictive features. These features are the first 45 PC maps derived from the training dataset and included in the prediction model and further projected to the MR-ART data normalized based on training data. While evaluating the effect of sessions we found significant difference after FDR correction between low motion session and no motion in 22 PCs (out of 45 PCs included, explaining 27% of the total variance in CamCAN data) and between high motion and no motion in 36 PCs (out of 45 PCs included, explaining 40% of the total variance in CamCAN data). The results for visually rated motion were very similar to the high motion session with 35 PCs showing significant relationship with motion levels after FDR correction (out of 45 PCs included, explaining 39% of the total variance in CamCAN data). The only differences between the two models were in PCs number 16, 23, and 10, The first two (i.e., PC-16 and PC-23 both showed significant difference between high and no motion but not a significant relationship with visual rating and PC-10 vice versa). We have shown the z-scored pattern of the first 4 PCs here as an example of the main patterns affected in both models.

**Figure 6.**
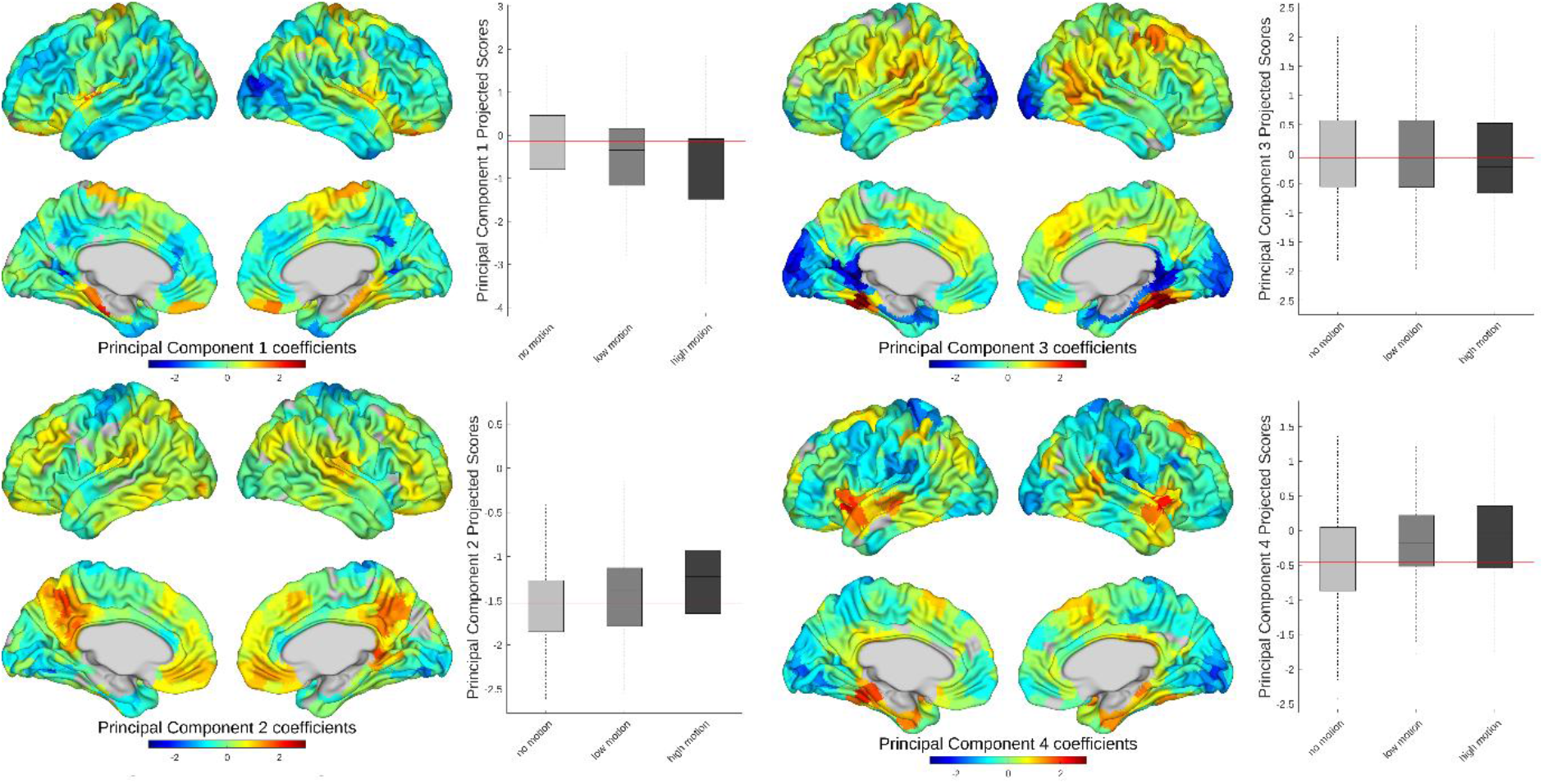
The normalized maps of the first 4 principal components used in age prediction. All four principal components are significantly different between no-motion and high-motion sessions after FDR correction. Each principal component map is accompanied by a boxplot for its corresponding scores for each MR-ART data point across three sessions.

## Discussion

In this work, we evaluated the influence of head motion on BAG estimation, determined by changes in brain age prediction derived from VBM and FreeSurfer cortical thickness measurements. Analyzing the independent MR-ART dataset, we observed that the same individuals rescanned with head motion induced during scanning exhibited significantly greater BAGs. Our study demonstrated a robust correlation between the prediction of BAG changes and the three motion levels of the MR-ART dataset. When compared to the session with no induced motion, the low-motion session exhibited a significantly higher brain age gap, with an average increase of 0.95 years (2.28 years for FreeSurfer-based results), while the high-motion session showed an even greater increase, with an average of 2.35 years (3.46 years for FreeSurfer-based results, see Table 3 for a summary of the ranges of disease-associated delta age reported in the literature). These results were further validated through motion level ratings assessed visually by an expert, where the results revealed a notable systematic change in the BAG, as evidenced by an average increase of 0.45 years per level of visually rated motion from 0 to 5 (0.83 years per level of visually rated motion for FreeSurfer-based results), approximately 1.5 to 2 years for high motion levels. Based on these findings we suggest addition of visual motion rating as an accompanying covariate in the BAG analysis especially for the population with higher reported motion. This step will enable the researcher to distinguish different sources contributing to BAG estimates (biological versus motion/symptom related changes). We further demonstrated that the normalized Euler number (based on formula (1)) can explain 70% of the variance in motion ratings. Based on this definition, each unit increase in the normalized Euler number is related to an average increase of 1.12 years increase in BAG per unit based on VBM as well as 2.5 years increase in BAG per unit based on cortical thickness. This is promising and suggests that for larger datasets as well as when trained experts for the visual motion rating are not available, this transformed Euler number can be used as a proxy measure.

**Table 3.**
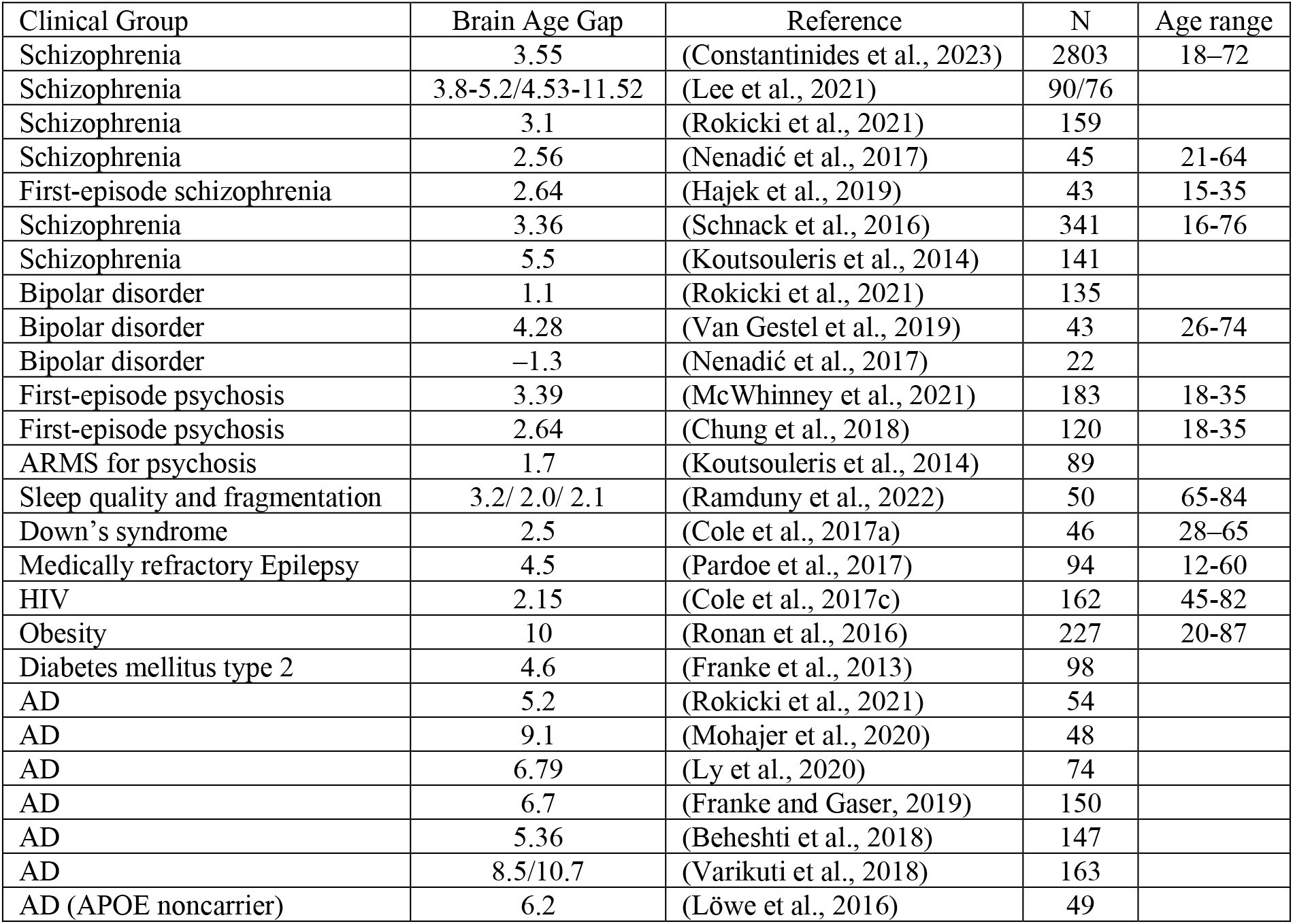
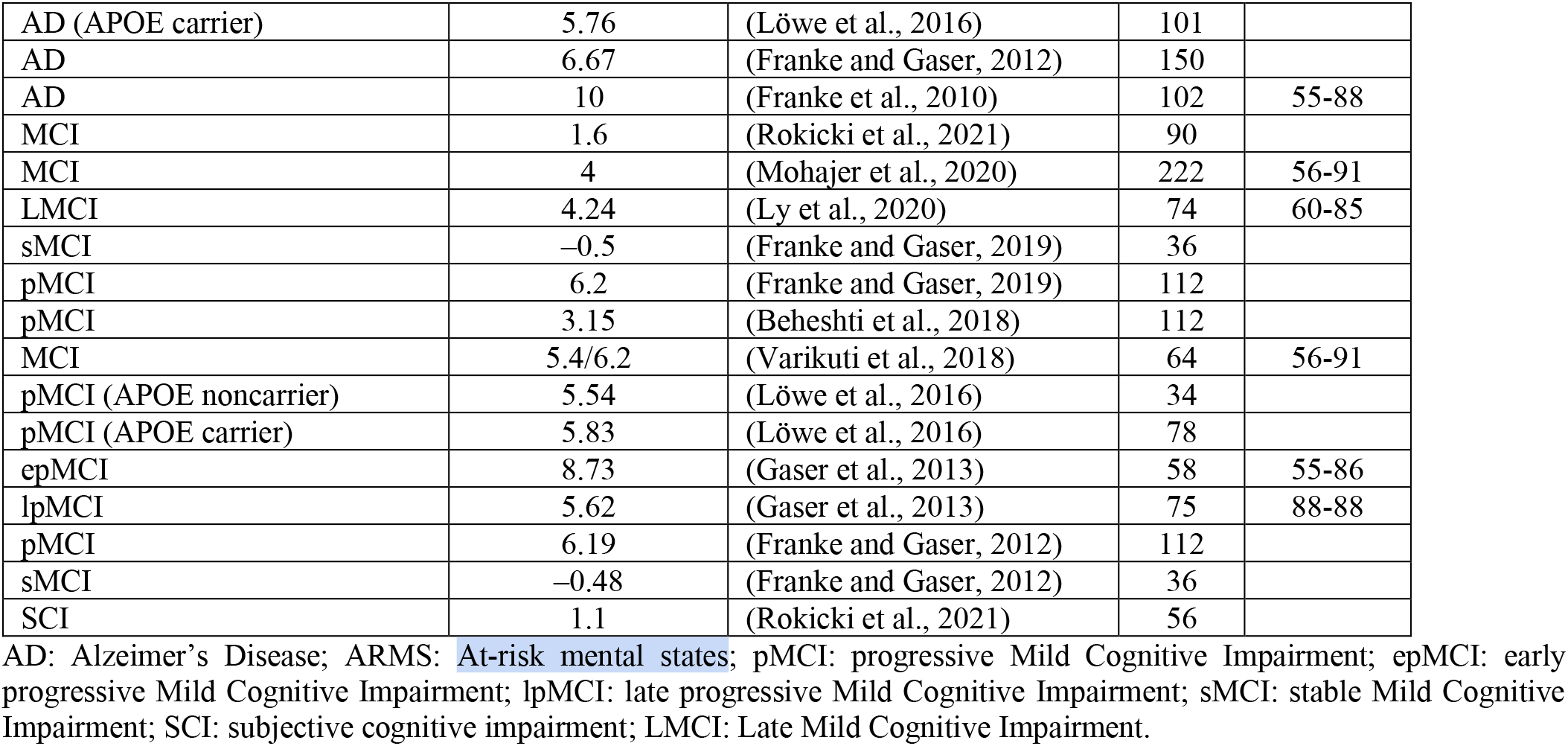
Brain age gap estimates in different disorders reported in the literature.

In line with previous studies in the literature (De Lange et al., 2022; Gao and Pang, 2022; Han et al., 2022; Monti et al., 2020), the brain age prediction model was trained based on a healthy aging population (CamCAN), and was then applied to the target MR-ART dataset, to provide an estimation of the range of effects sizes of delta brain age values that can be attributed to motion. As a complementary step we also used the MR-ART dataset for both training and test, where we trained the model on the data from no motion session and tested it on high motion session and vice versa. Based on this analysis, while it is expected that the models would perform better when the training and testing data are more similar, this source of variability would have resulted only in higher variance in the prediction for the test group. However, our results identify a directional bias in the prediction in both sets of analysis showing lower BAG in no motion session compared to high motion session. This clearly suggests that higher motion will bias the prediction towards higher BAG (i.e., older brain age) in a biased manner as opposed to only adding variance in the prediction model which will inflate the findings from study of BAG in participants with various disorders.

BAG is a widely used measure of brain health that allows for assessment of individual age-related brain changes, providing a personalized biomarker of brain health (Cole et al., 2017b; Franke et al., 2014, 2010; Gaser et al., 2013; Sendi et al., 2021). BAG can be potentially used to determine biological aging, predict rate of aging, and to monitor the fundamental processes of aging non-invasively (Gaser et al., 2013; Sendi et al., 2021). Furthermore, BAG can be used to assess and quantify accelerations in brain changes due to various brain disorders (Franke et al., 2014). Hence, BAG provides a more practical approach for assessing and tracking individual brain aging trajectories and its deviation from normative brain aging over time (Franke et al., 2010). As such, numerous recent studies have employed BAG estimates as biomarkers of brain health (Franke and Gaser, 2019).

As a biomarker, BAG has provided a reference for age-related alterations in brain structure. Moreover, it has demonstrated significant potential for convenient implementation in multi-center studies. The predictive accuracy obtained from assessing the structural and functional brain age could be instrumental in tailoring personalized treatments and interventions (Franke and Gaser, 2019). It has been shown that late-life cognitive decline can be predicted in patients with mild cognitive impairment and AD based on the brain age gap (Franke and Gaser, 2019). Furthermore, research indicates that a higher brain age gap during midlife is preceded by a decline in cognitive functioning that begins in childhood and continues over time (Elliott et al., 2021). Although it may be intuitive to associate a higher brain age gap with unhealthy lifestyle choices, such as smoking or alcohol consumption, and subsequent cognitive decline, our findings indicate that the predicted increase in BAG can also be influenced by motion artifacts (Elliott et al., 2021; Franke and Gaser, 2012).

The impact of head motion on structural and functional brain measure estimations has been well-documented in previous studies. Head motion, a common occurrence during MRI scanning, has the potential to significantly affect MRI data and act as a major confounding factor in MRI-derived models. These effects include but are not limited to a reduction in gray matter volume as well as blurring between the cortical gray and white matter boundaries and consequently change in cortical thickness (Alexander-Bloch et al., 2016; Bedford et al., 2020; Pardoe et al., 2016; Reuter et al., 2015). Given the negative association of age with brain volume and cortical thickness, both of these effects will result in higher age estimation and consequently higher BAGs. During adulthood (Madan, 2018; Pardoe et al., 2016; Pollak et al., 2023), motion levels are also higher in older populations and particularly in individuals with obesity (Beyer et al., 2020; Zeighami et al., 2021), PD (Makowski et al., 2019; Torres and Denisova, 2016), and other neurodegenerative disorders. Therefore, systematic biases in the estimated BAGs in these populations can confound the biologically relevant portions of BAG. Therefore, it is crucial to address and account for head motion effects when interpreting age-related differences in brain measures.

In a multimodal brain age prediction study,(Liem et al., 2017) examined the effect of motion as measured based on functional MRI, and concluded that head motion did not drive brain-based age prediction. However, the study also reported that the removal of motion significantly increased the absolute prediction error for brain age, suggesting that part of the brain age prediction was driven by the motion. There are a number of major differences between the present study and the approach by Liem et al. 2016. Our study mainly focuses on structural MRI motion, whereas Leim et al. assesses motion based on functional MRI. Furthermore, as an initial step, Liem et al. removed scans with excessive motion as defined by a mean Framewise Displacement (FD) exceeding 0.6 mm for functional scans which is significantly lower than the ranges reported in studies of patient populations (e.g. 1-1.2 mm and 1-2 mm for Parkinson’s disease and stroke patients, respectively) (Guo et al., 2022; Seto et al., 2001). This can partially explain the study’s conclusion with regards to motion. A threshold of 0.6 mm mean FD would however be a very conservative exclusion criteria for structural MRI studies due to their robustness to motion and prove prohibitive to studying the more severe cases of the disorders. Another step to infer that motion effect was not a confound in their study was to motion matching and generalizability as the confirmation of robustness of findings against motion. However, in this step as well the study has used “a motion adjusted subsample of the test set, created by restricting the sample to subjects with a mean FD between 0.19 and 0.28 mm” which can potentially limit the generalizability of the conclusion and explain the differences in our findings.

In several studies which directly utilize structural MRI to predict brain age, the Euler number, a measure calculated by FreeSurfer as a proxy for MRI processing quality has been used either as an exclusion criteria (Madan, 2018) or as a covariate (alongside age and sex) in examining the BAG differences between groups. We believe that these approaches, especially using Euler number as a covariate will remove a portion of motion related signal. However, based on our results from mixed effect modeling, normalized Euler number explains 60% of the variance in our motion rating (compared to 30% variance explained by raw Euler number) and therefore it will be a more effective covariate in the future modeling approaches, at least with regards to motion. Finally, in comparison with the previous studies, our results provide a systematic approach to quantify these effects as well as an estimate for the effect size of this given bias for a given motion level. Furthermore, we have made our motion ratings for the open access MR-ART datasets publicly available (Supplementary Table 1), which can help other researchers estimate and remove motion-related confounding factors from their studies.

While presence and severity of motion in the scanner can be related to presence of neurological disorders such as Parkinson’s disease and a motion-related increase in brain age might also imply greater disease severity, the underlying assumption of brain age prediction models is that they provide an estimation of brain health based on its underlying biology as measured by MRI, and as such, higher levels of estimated motion that are caused by errors in the estimated grey matter features (as opposed to real change in gray matter density or cortical thickness) are not desirable. MR-ARTs provided us with the unique opportunity to model a realistic signal which is decorrelated from its natural source as induced motion in the MR-ARTs was similar to the signal observed in patient cohorts (e.g., subjects with obesity or Parkinson’s disease, see Supplementary Figure S.2 for an example) in terms of pattern and severity, but was not related to presence of any disorders.

Our study benefits from several strengths that enhance the robustness and reliability of our findings. Firstly, we conducted training and validation on different datasets, ensuring the generalizability of our results beyond the specific dataset used for training. This approach enhances the validity of our conclusions and allows for a more comprehensive assessment of the predictive accuracy of our models. Secondly, our study used a validation dataset comprising scans from the same individuals scanned three times with different motion levels, enabling direct comparisons of brain age estimates based on motion within the same subject. This internal validation provides a rigorous assessment of our models and preprocessing methods, adding further credibility to our findings. Furthermore, we have used both VBM (using in-house pipelines) and cortical thickness (as measured by FreeSurfer 7.4.1) two widely used measures in the study of brain aging, neurodegeneration, and BAG, and identified the presence of the effect in both analysis, suggesting that the results are not dependent on the particular measure or software used. The results are also similar when we used multiple predictive models including SVM with linear Kernels, Gaussian Process predictive modeling, as well as tree-based approaches which suggest the effect of motion in BAG is not dependent on the machine learning methods used.

The image acquisition method in MR-ART datasets was designed to encompass three levels of motion for each subject, ranging from no motion to a high level of motion. It is important to note that the levels of motion were closely related to human actions; hence, the three motion labels did not precisely represent distinct categories within these specified levels of motion. Consequently, during expert evaluation, the datasets were effectively categorized into six distinct levels of motion, to provide more accurate measurements of the severity of motion occurring in each image, as not all participants had exhibited the same level of motion in each session. BAGs showed an even stronger relationship with motion severity ratings, further supporting our results.

Our study is not without limitations. Firstly, our sample from MR-ART consisted only of healthy participants from a relatively limited age range, which may limit the generalizability of our findings to more vulnerable population groups or people in later life stages. Utilizing more diverse and heterogeneous samples could enhance our ability to identify the impact of motion on the BAG. Secondly, the motion dataset did not include behavioral measures that could have provided valuable insights into differences in the relationships between motion-impacted BAG and cognitive scores. The absence of such data restricts our ability to fully explore the potential links between motion artifacts and the brain estimates of cognitive outcomes. Additionally, the motion dataset may not be fully representative of natural motion occurring in the scanner or motion associated with specific diseases. The patterns of motion observed in the dataset might not entirely mirror the natural motion experienced by individuals during brain imaging scans, which can introduce potential biases in our analysis. Furthermore, here we are using a rather simple model for brain age prediction. While we compared our model with other methods such as support vector machines and found very similar results and no change in performance or conclusions, future studies using more advanced methods such as deep neural network models for age prediction might further improve the prediction performance and the motion bias estimation. Finally, future studies are needed to simulate natural motion artifacts and provide solutions for robust motion removal that can recover the brain age estimates based on the no motion data from these motion contaminated scans.

While the training and test datasets were acquired using different scanner models, the impact of motion was assessed by comparing the model estimations of the same individuals scanned using the same scanner, without motion or with different levels of motion. Given the accuracy of the brain age prediction model for the scans that did not include any motion, as well as the fact that the same scanner was used to acquire the motion induced images, it is unlikely that the results would be impacted by scanner differences between the train and test datasets. Finally, in the validation analyses (Section 3.3), the model was trained based on the same (MR-ART) dataset, yielding similar results, suggesting that the findings are not due to scanner related differences between training and test datasets.

In conclusion, we showed that participants’ motion in the scanner during acquisition of T1w images can significantly and systematically impact the derived BAG estimates. Our results suggest that brain age studies would benefit from considering assessment of motion severity, particularly visual rating of the input scans and potentially including motion severity as a covariate in the models.

## Data and Code Availability

CamCAN and MR-ART datasets are available at https://www.cam-can.org/index.php?content=dataset and https://openneuro.org/datasets/ds004173/versions/1.0.2. We have also made our motion severity ratings available as a Table (Table S.1) in the Supplementary Materials. The scripts for MRI data preprocessing and VBM calculations are available at https://github.com/NIST-MNI/nist_mni_pipelines.

## Author Contributions

Roqaie Moqadam: Motion ratings, analyses, manuscript writing.

Mahsa Dadar: Study design, analyses, manuscript writing and revision.

Yashar Zeighami: Study design, analyses, manuscript writing and revision.

## Acknowledgements

The training dataset was obtained from the Cambridge Centre for Ageing and Neuroscience (CamCAN). CamCAN funding was provided by the UK Biotechnology and Biological Sciences Research Council (grant number BB/H008217/1), together with support from the UK Medical Research Council and University of Cambridge, UK. HCP data was obtained from the Human Connectome Project, WU-Minn Consortium (Principal Investigators: David Van Essen and Kamil Ugurbil; 1U54MH091657) funded by the 16 NIH Institutes and Centers that support the NIH Blueprint for Neuroscience Research; and by the McDonnell Centre for Systems Neuroscience at Washington University.

Authors also acknowledge Compute Canada (https://www.computecanada.ca/home) for the usage of the computing resources in the current work.

